# Evaluating Synthetic Diffusion MRI Maps created with Diffusion Denoising Probabilistic Models

**DOI:** 10.1101/2024.11.06.621173

**Authors:** Tamoghna Chattopadhyay, Chirag Jagad, Rudransh Kush, Vraj Dharmesh Desai, Sophia I. Thomopoulos, Julio E. Villalón-Reina, Paul M. Thompson

## Abstract

Generative AI models, such as Stable Diffusion, DALL-E, and MidJourney, have recently gained widespread attention as they can generate high-quality synthetic images by learning the distribution of complex, high-dimensional image data. These models are now being adapted for medical and neuroimaging data, where AI-based tasks such as diagnostic classification and predictive modeling typically use deep learning methods, such as convolutional neural networks (CNNs) and vision transformers (ViTs), with interpretability enhancements. In our study, we trained latent diffusion models (LDM) and denoising diffusion probabilistic models (DDPM) specifically to generate synthetic diffusion tensor imaging (DTI) maps. We developed models that generate synthetic DTI maps of mean diffusivity by training on real 3D DTI scans, and evaluating realism and diversity of the synthetic data using maximum mean discrepancy (MMD) and multi-scale structural similarity index (MS-SSIM). We also assess the performance of a 3D CNN-based sex classifier, by training on combinations of real and synthetic DTIs, to check if performance improved when adding the synthetic scans during training, as a form of data augmentation. Our approach efficiently produces realistic and diverse synthetic data, potentially helping to create interpretable AI-driven maps for neuroscience research and clinical diagnostics.

## 1. INTRODUCTION

In recent years, the intersection of deep learning and biomedical imaging has opened new avenues to generate synthetic images, with significant implications for research and clinical practice. Neuroimaging techniques, such as magnetic resonance imaging (MRI) and and diffusion tensor imaging (DTI), can provide detailed insights into the brain’s gross anatomy and white matter microstructure, but acquiring large datasets can be challenging due to the high cost, time constraints, and the need for patient consent for their images to be used in research. These obstacles often hinder the development and validation of robust diagnostic tools and data-driven treatments. Synthetic images generated through deep learning models offer a possible solution to these limitations. By creating realistic neuroimaging data, researchers can augment existing datasets, making it easier to develop and validate new diagnostic tools and even simulate the effects of treatment strategies. Deep learning has made it possible to generate synthetic neuroimaging data through the use of generative models such as Generative Adversarial Networks (GANs) [1] and Variational Autoencoders (VAEs) [2]. GANs’ adversarial training process can produce highly realistic images, yet they can be unstable and difficult to train. Common issues with GANs include mode collapse, where the generator produces a limited variety of images, and sensitivity to hyperparameters, which can lead to inconsistent output quality. VAEs, by contrast, learn a latent representation of the data by encoding images into a latent space and then decoding them back to their original form. While VAEs are generally more stable and easier to train, they often suffer from blurry reconstructions, which limits their effectiveness in generating high-resolution medical imaging data.

To address these challenges, researchers have increasingly turned to denoising diffusion probabilistic models [3], which offer a robust alternative for generating high-quality synthetic images. Unlike GANs and VAEs, denoising diffusion models gradually transform a simple noise distribution into a complex data distribution through a series of iterative refinements. These models are trained to reverse a noising process, effectively learning to generate data by progressively denoising random noise until a coherent image emerges. This approach can produce more diverse and high-fidelity images, while also being more stable to train compared to GANs. This type of image generation has also been combined with conditioning approaches, where learned embeddings are used to condition images based on a text prompt (as in stable diffusion [3], or based on classifier-free guidance [4], to generate images that match a specific class. In the context of neuroimaging, denoising diffusion models can generate synthetic images that preserve the fine anatomical details necessary for quantitative analysis but also overcome the common issues of blurriness and mode collapse associated with other generative models. Pinaya et al. [5] developed latent diffusion models (LDM) to produce realistic brain MRI scans, conditioning the generation process on demographic factors such as age and sex. Additionally, various forms of DDPMs have been employed by other researchers to create synthetic brain MRIs [6,7,8,9]. Diffusion modeling techniques can also be applied to counterfactual image generation [10,11,12], where individual images are modified to align with a reference distribution derived from healthy controls, while preserving the unique anatomical characteristics of the subject. They have also been applied to cross-modality image translation [13]. Most of these DDPM models focus on T1-weighted MRI, CT scans or PET scans, usually of the human brain. In this work, we focus on training diffusion models to generate synthetic diffusion-weighted images, specifically, DTI-MD images - a scalar map commonly generated from diffusion-weighted MRI, which are valuable for characterizing white matter microstructure and are widely used in clinical and research settings. We assessed methods to train diffusion models under constraints of limited data, computational time, and memory resources. This involved systematically adjusting the model size, training duration, and incorporating a latent diffusion model to optimize performance. Moreover, we evaluate the utility of these synthetic datasets for data augmentation in downstream tasks –such as sex classification using convolutional neural networks (CNNs)--providing a benchmark for their practical applications in data-driven neuroscience.

## 2. DATA AND PREPROCESSING

Diffusion tensor imaging (DTI) is the most commonly used technique for modeling brain tissue microstructure *in vivo*. A variant of standard MRI, diffusion-weighting is used to sensitize the MRI signal to the rate of water diffusion in a range of spherical directions. The standard diffusion tensor model approximates the local diffusion process using a spatially-varying diffusion tensor, typically represented as a 3D Gaussian approximation at each image voxel. While the tensor model simplifies the diffusion process, making it less complex than higher-order models such as NODDI [14] and MAP-MRI [15], it remains widely used due to its compatibility with single-shell diffusion MRI (dMRI), which is faster and more convenient for clinical applications. DTI is typically summarized using four standard scalar metrics: fractional anisotropy (FA), and mean, axial, and radial diffusivity (MD, AxD, and RD). These metrics describe the shape of the diffusion tensor at each voxel, derived from the three principal eigenvalues of the tensor, which represent the rate of water diffusion in 3 principal directions (eigenvectors) at each point in the tissue. Among these measures, MD is particularly sensitive to changes in cellularity, edema, and necrosis, and is among the most sensitive measures to neurodegenerative diseases such as Alzheimer’s disease and Parkinson’s disease [16,17].

The standard formula for MD, in terms of the diffusion tensor eigenvalues, is:

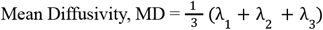

The primary dataset for our experiments is the Cambridge Centre for Aging and Neuroscience (Cam-CAN) dataset [18], which we selected as it spans a larger proportion of the human lifespan than many other datasets. We used data from a total of 652 healthy control (CN) participants (mean age: 54.29 ±18.59 (SD) years; 322 F/330 M). The dMRI processing pipeline that we used is extensively detailed in [19,20]. All diffusion models were trained on 3D DTI-MD maps from Cam-CAN.

## 3. METHODS

Diffusion models are generative in nature, approximating the underlying data distribution through a distribution that is learned by iterative approximation, using a trained neural network. During the training phase, noise sampled from a normal distribution is progressively added to an image according to a noise scheduler, such as the linear scheduler, where noise is introduced at discrete steps. Models such as the Diffusion Pixel-Space based [5] DDPM and Latent Diffusion Models (LDM) typically employ a modified U-Net architecture enhanced with a self-attention mechanism. The discrete time steps can also be used to condition the model, using a cross-attention mechanism that is integrated into the ResNet blocks within the architecture. The primary objective of the diffusion model is to predict the noise added at a random time step, which is then used to minimize a regression loss function, commonly mean squared error (MSE). Once trained, the model can generate synthetic scans through different schedulers, such as the DDPM or its deterministic variant, the Denoising Diffusion Implicit Model (DDIM) [5], with a predefined number of sampling steps. During sampling, the model iteratively transforms pure noise into an image that is drawn from a distribution that closely approximates the training distribution. The choice of sampling scheduler plays a crucial role in determining the quality of the synthetic scans and the time required to generate each scan. The conditional version of the DDPM can be created, adapting the original model to incorporate a cross-attention mechanism, enabling external conditioning through classifier-free guidance, or even using a text prompt, which can be jointly embedded with the image using models based on CLIP [21]. This modification allows the model to generate images based on additional information, such as the sex, age, or diagnosis of the subject. Unlike DDPMs, LDMs use autoencoders to compress images into a latent space before applying standard diffusion modeling. This approach helps to mitigate the computational challenges associated with training large deep learning models on 3D MRI scans, where memory requirements can be high. The autoencoder in LDMs is trained using a combination of losses, including a reconstruction loss, perceptual loss, KL divergence loss, and an adversarial loss [3,4].We trained one version of each LDM and DDPM architecture.

As is standard in the evaluation of generative models, we quantitatively assessed our models for realism and diversity of the resulting synthetic images using performance metrics. Metrics were averaged over 500 evaluation pairs, with the Maximum Mean Discrepancy (MMD) [22] and Multi-scale Structural Similarity Index (MS-SSIM) to measure similarity. Lower MMD values indicate closer distributions (meaning that the synthetic data closely resembles the real data distribution), while higher MS-SSIM values denote greater similarity. For realism, MMD and MS-SSIM were used to compare real and synthetic images of the same class. Diversity was evaluated by comparing two unique synthetic images conditioned on the same class using MS-SSIM, with MS-SSIM also applied to real image pairs as a reference.

The 3D CNN architecture (**Fig. 3**) consisted of four 3D Convolution layers with a 3×3 filter size, followed by one 3D Convolution layer with a 1×1 filter, and a final Dense layer. All layers used the ReLu activation function and Instance Normalization. Dropout layers, with a dropout rate of 0.5, and a 3D Average Pooling layer with a 2×2 filter size were added to the 2^nd^, 3^rd^, and 4^th^ layers. Models were trained with a learning rate of 10^−4^, and test performance was assessed using balanced accuracy. We trained this CNN model for 100 epochs, with a batch size of 8. The learning rate was exponentially decayed with a decay rate of 0.96. The Adam optimizer [23] and mean square error loss function were used for training. To deal with overfitting, dropout between layers and early stopping were used. Test performance on Sex Classification was assessed using the Balanced Accuracy and F1 Score to compare results for different experiments. After registering the images to a common template, the data was split into independent training, validation, and testing sets in the ratio of approximately 70:20:10. Two different downstream experiments were conducted - first, starting with all of the real images for training and systematically adding synthetic images while decreasing the proportion of real images; and second, systematically adding synthetic images while keeping the number of real images in training constant. These experiments were conducted for all models - DDPM, Conditional DDPM, LDM and Conditional LDM.

**Figure 1.**
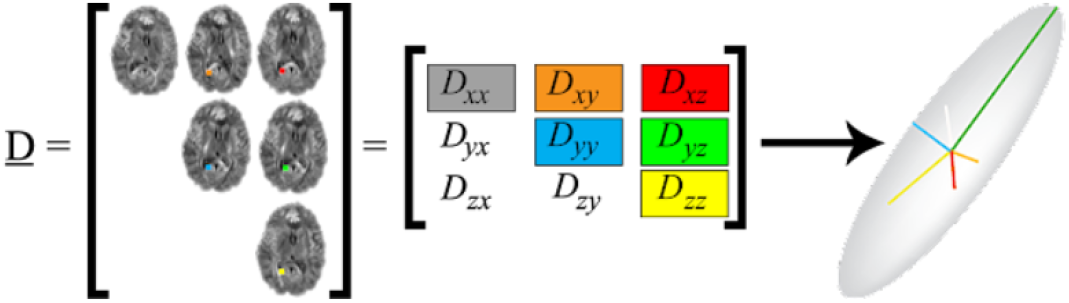
Diffusion Tensor Components - In the simplest case, the diffusion tensor represents a 3D Gaussian model of water diffusion, modeled using a diffusion ellipsoid, or a 3×3 matrix representing diffusion rates in different directions; this tensor can be rotated to assume a diagonal matrix form with only 3 diagonal components - the eigenvalues - which represent the relative diffusion tendencies in each of 3 principal directions.

**Figure 2.**
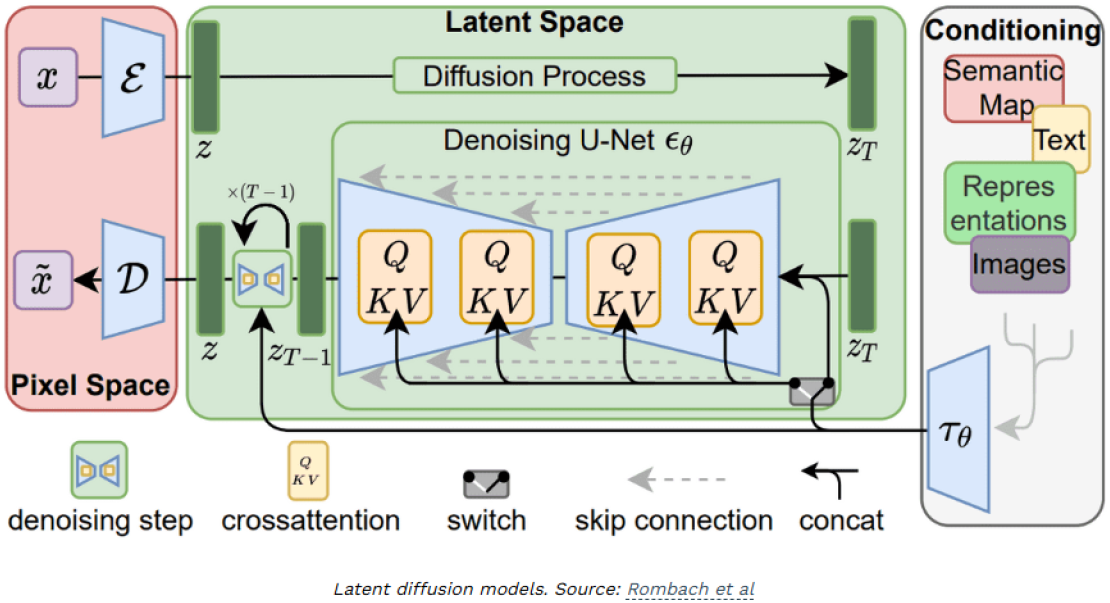
Latent Diffusion Model Architecture, reproduced from Rombach et al. [3].

**Figure 3.**
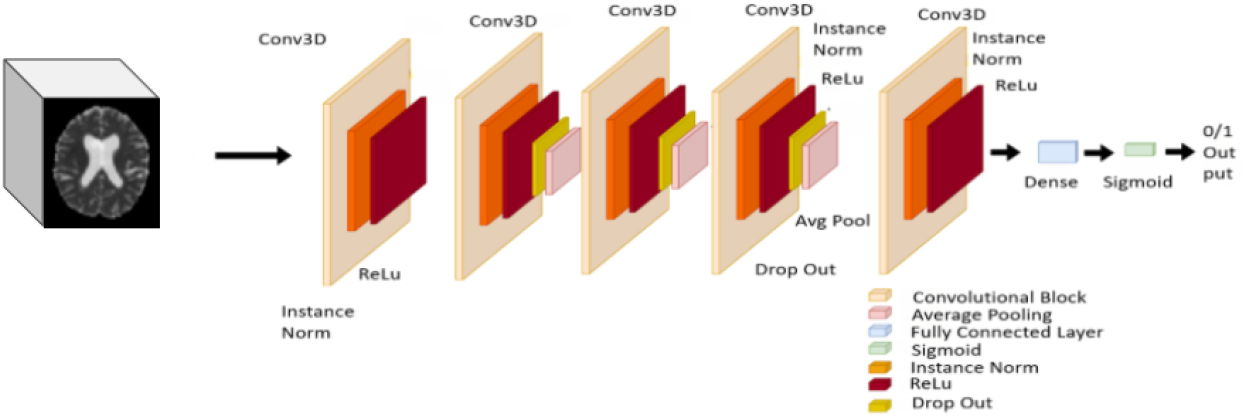
3D CNN Architecture that we trained for downstream tasks.

**Figure 4.**
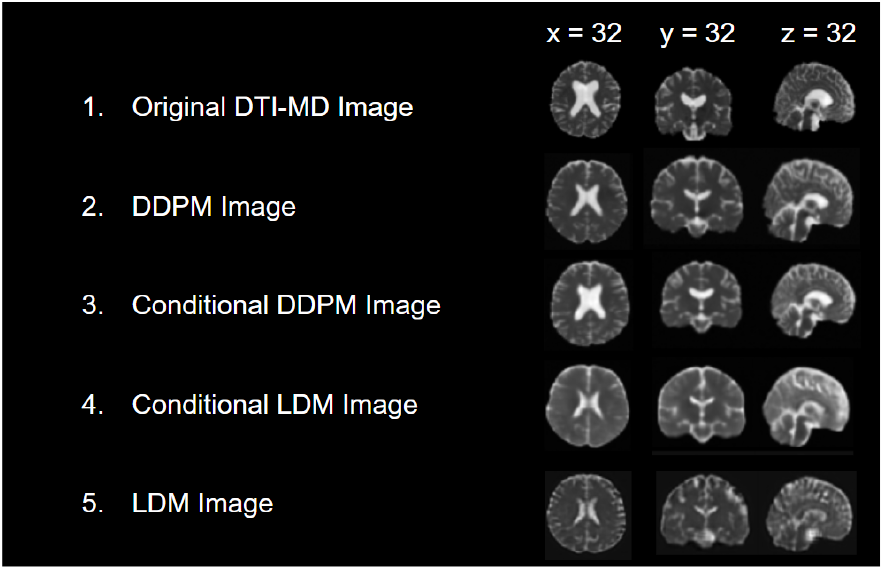
The first row shows the original real DTI-MD maps from an example subject in the Cam-CAN dataset. The next four rows show the Synthetic DTI-MD maps generated by the various models - DDPM, Conditional DDPM, Conditional LDM and LDM. Notably, some of the sulcal features (fissures) in the cortex are visible, even though the input data is only 2mm isotropic (a commonly used spatial resolution for DTI). The conditional variable is sex of the person - male or female. The slices in figure are all from male subjects.

**Figure 5.**
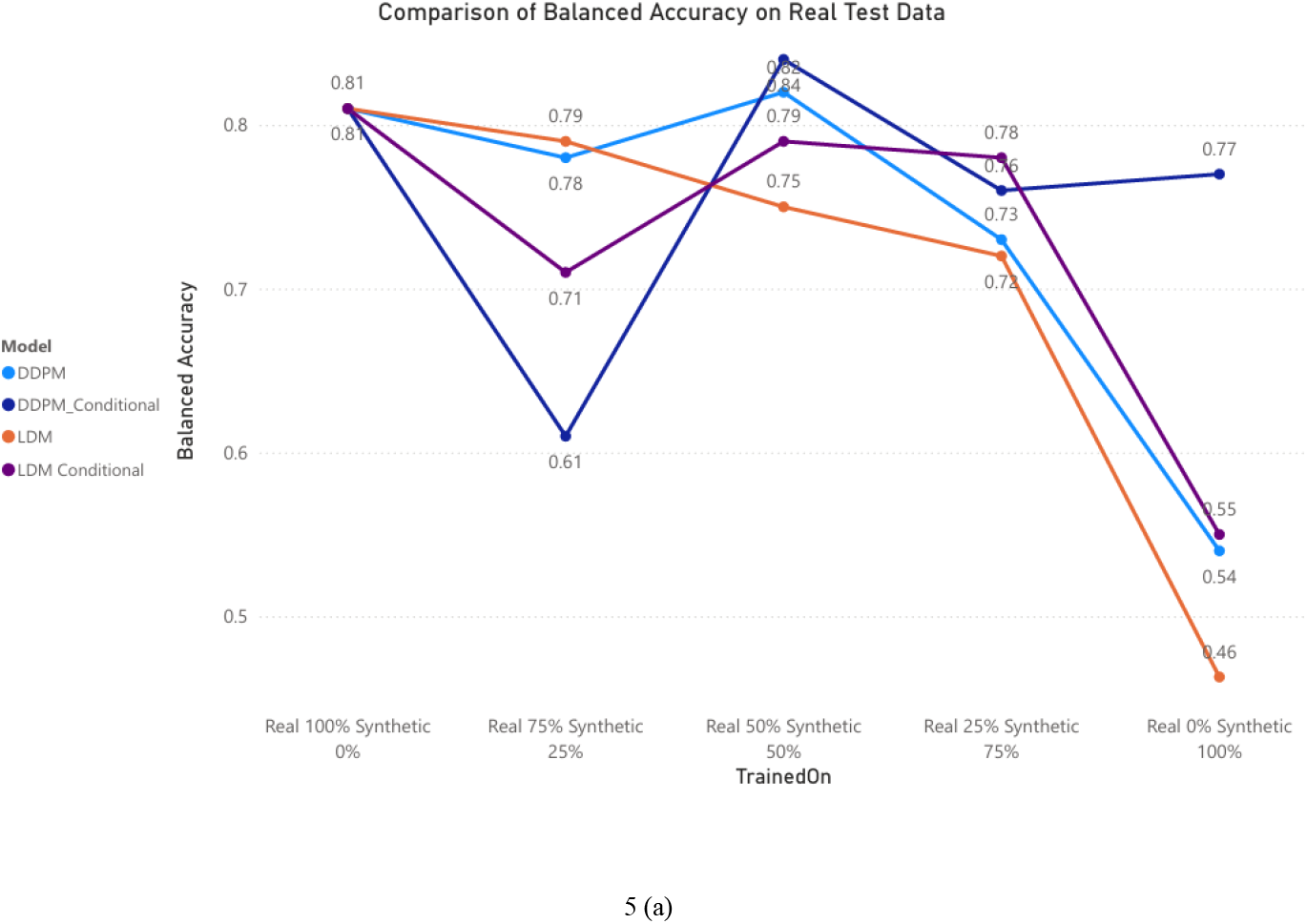
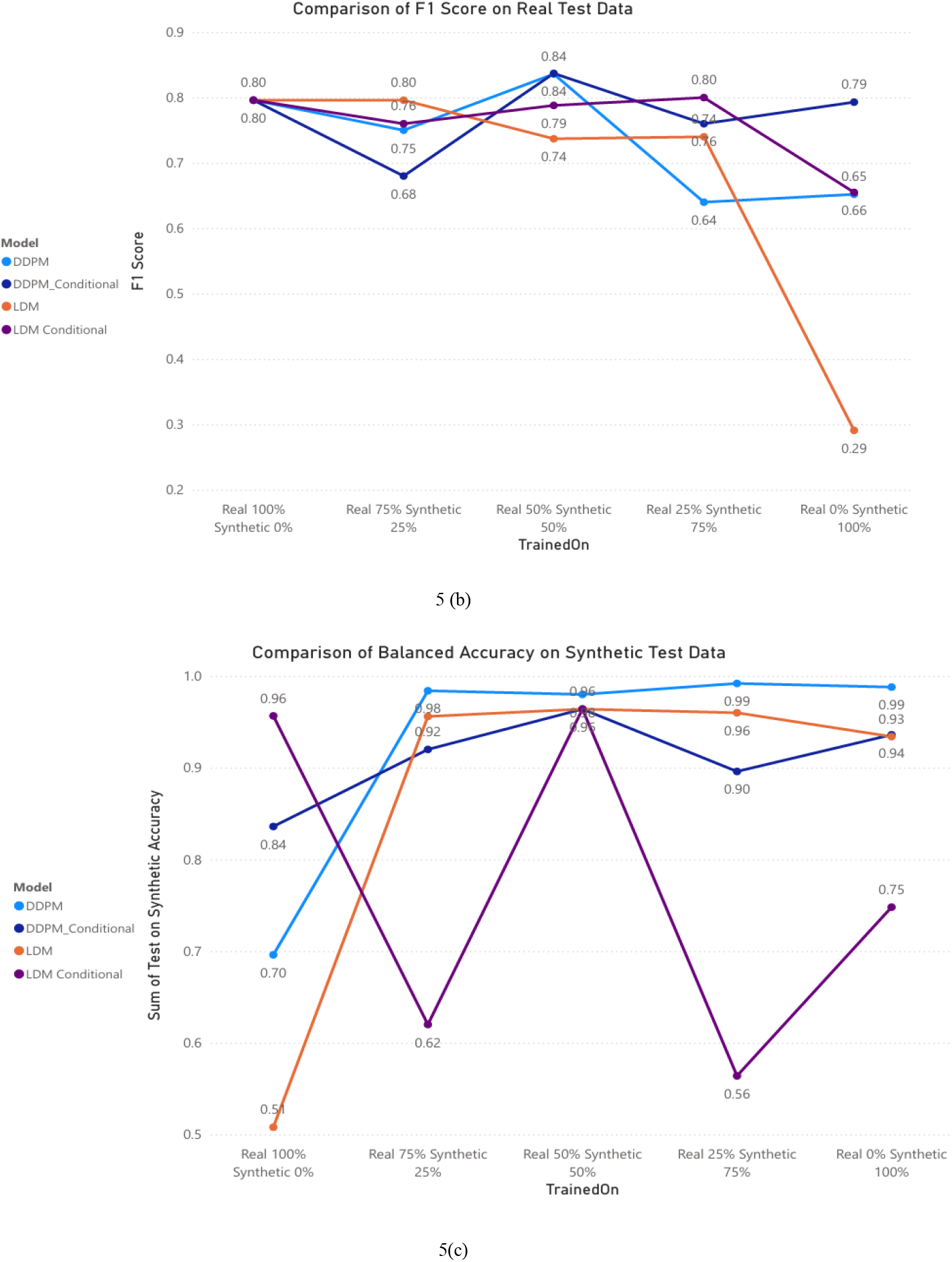
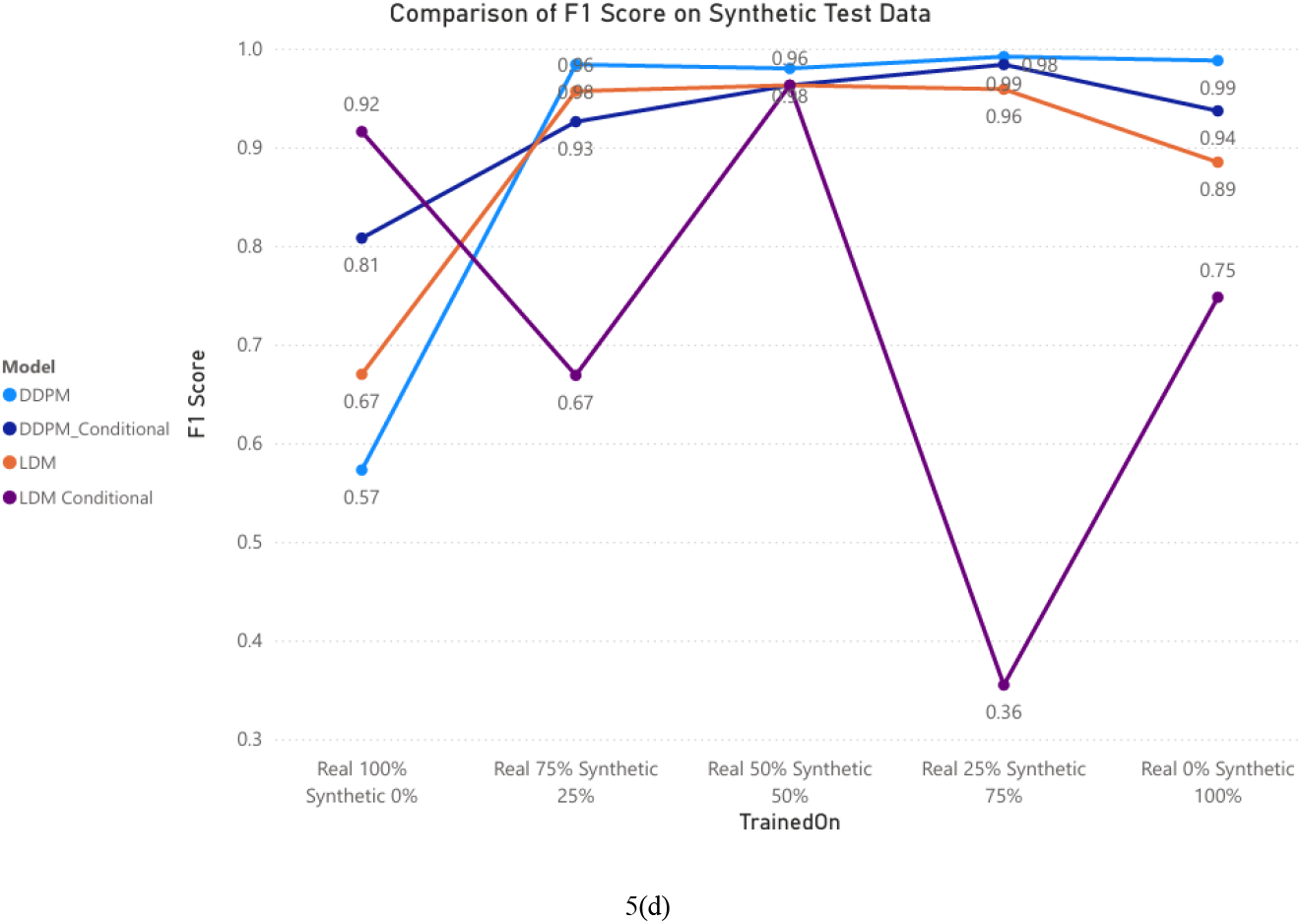
5(a) Comparison of Balanced Accuracy on Real Test Data. 5(b) Comparison of F1 Score on Real Test Data. 5(c) Comparison of Balanced Accuracy on Synthetic Test Data. 5(d) Comparison of F1 Score on Synthetic Test Data.

**Figure 6.**
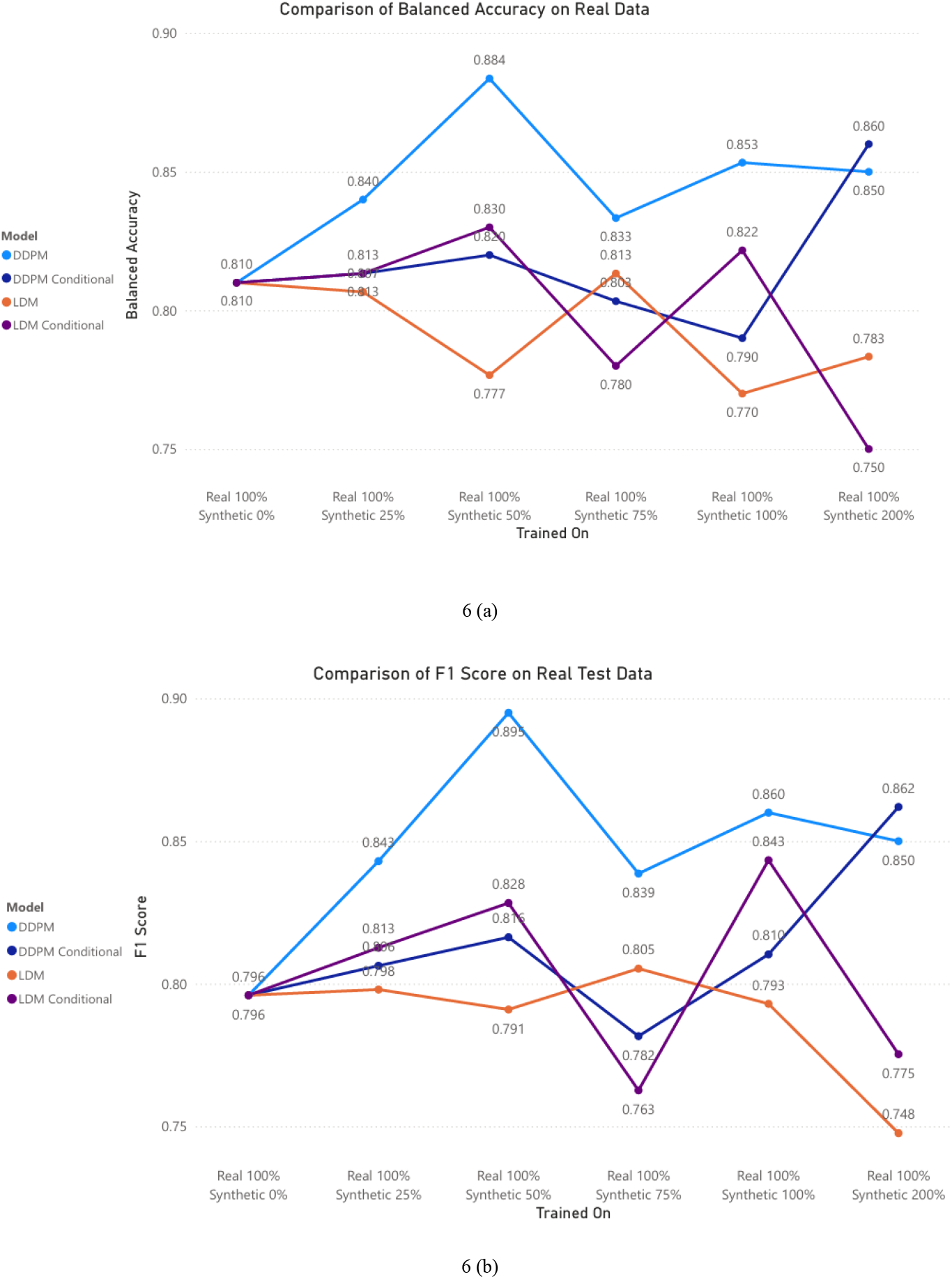
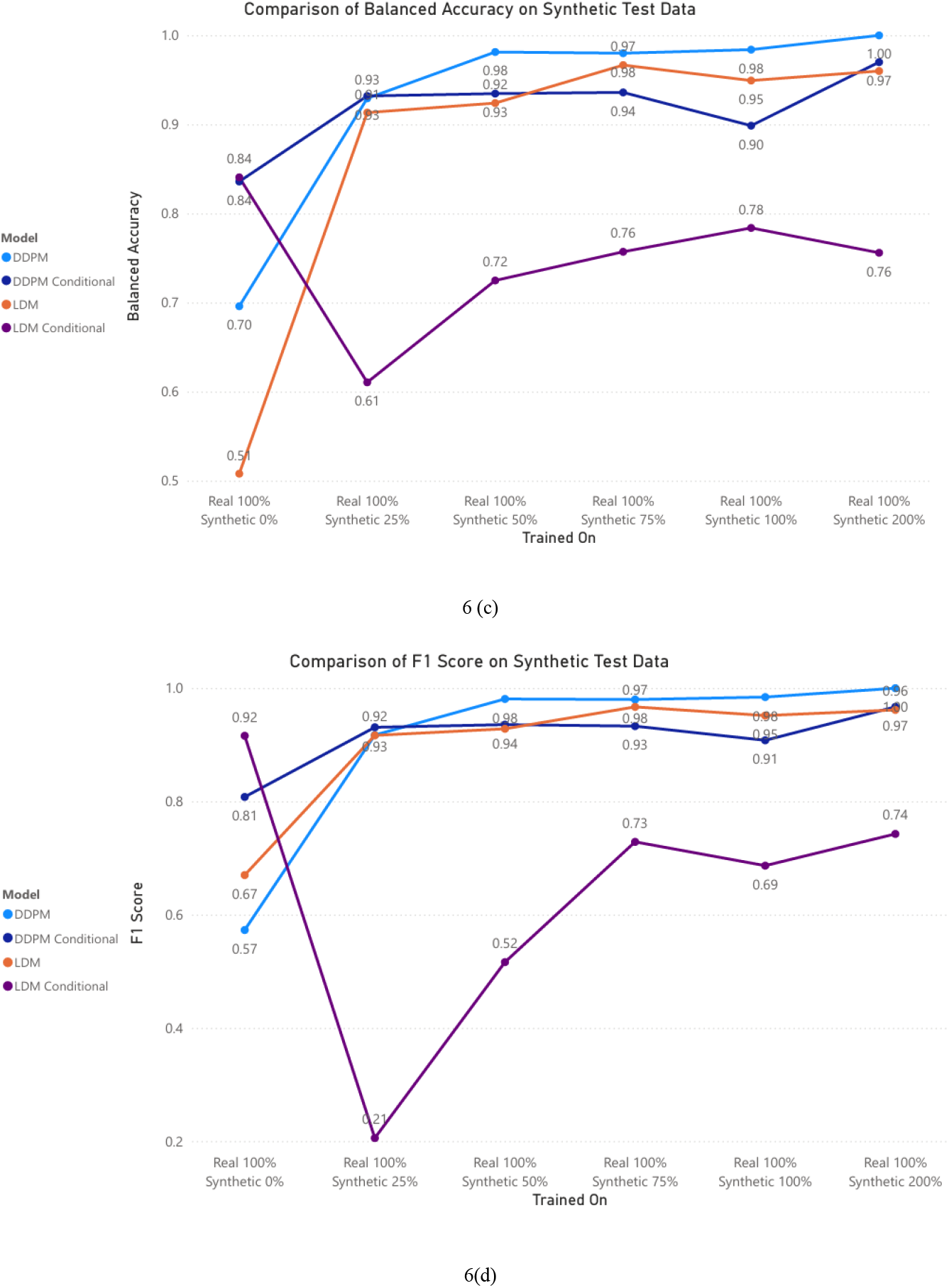
6(a) Comparison of Balanced Accuracy on Real Test Data. 6(b) Comparison of F1 Score on Real Test Data. 6(c) Comparison of Balanced Accuracy on Synthetic Test Data. 6(d) Comparison of F1 Score on Synthetic Test Data.

## 4. RESULTS AND DISCUSSION

Evaluations of the synthetic DTI-MD maps generated by the different models as compared to the real scans are summarized in **Table 1**. The best performances are in bold. The results show that for both male and female subjects, LDM models show better performance on Realism Metrics, whereas DDPM models show better performance on Diversity Metric of Synthetic vs Synthetic. The diversity metric scores are fairly high, but it was expected given the high scores for Real vs Real images as well, which offer a reasonable lower bound for what can be achieved for this metric. One reason for this could be the use of the latent space in the LDM architecture, resulting in more similar generated images. The enhanced realism of LDMs likely stems from their ability to model fine-grained features more effectively, possibly due to architectural differences or superior latent representation learning. Conversely, the stronger diversity in DDPM-generated samples may be attributed to their broader sampling distributions or reduced reliance on restrictive priors, which can facilitate greater variability in outputs. The interplay between realism and diversity underscores the importance of balancing these metrics in evaluating generative models, as improvements in one aspect may influence the other.

**Table 1.**
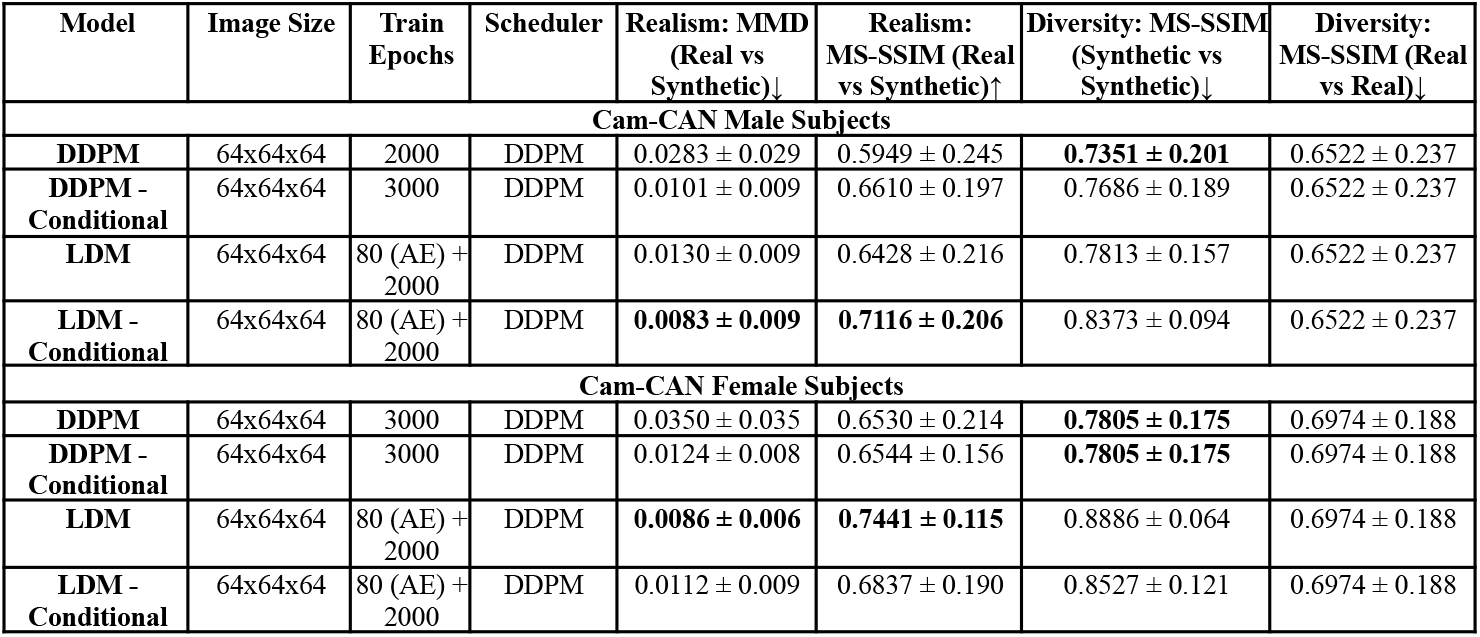
Realism and Diversity Test Metrics for the DDPMs and the LDMs. The best model performance for each metric is shown in bold. The MMD is computed from pairs drawn from the real and synthetic data distributions and lower values are better. SSIM is also a measure of realism, where higher values are better when real and synthetic images pairs are compared. SSIM can be adapted to measure diversity in synthetic images by interpreting low similarity as high diversity. A high SSIM value (close to 1) indicates that two images are very similar, whereas a lower SSIM value indicates greater dissimilarity. To use SSIM for measuring diversity, you would consider lower SSIM scores as indicative of greater diversity between pairs of images. In the last two columns, the diversity of the real data is considered to be a reasonable lower bound for this metric.

For the first downstream experiment, we evaluated the performance of CNN models trained on the real and synthetic DTI-MD maps for sex classification. The balanced accuracy and F1 scores were assessed on both real and synthetic test datasets across four models: DDPM, DDPM_Conditional, LDM, and LDM_Conditional. The experiment was designed such that in progressive runs, the amount of synthetic data in the training dataset was reduced by 25% and was replaced by real data from the Cam-CAN dataset. This was done until in the final run, the model was training on only real data. From the balanced accuracy on real test data, DDPM_Conditional consistently performed the best, achieving a peak balanced accuracy of 0.84 when trained with 50% real and 50% synthetic training data. This suggests that incorporating a small fraction of synthetic data does not degrade performance and may help the model generalize. The LDM models underperformed compared to the DDPM variants. This trend is echoed in F1 scores on real data, where DDPM_Conditional outperformed other models, while LDM models struggled, particularly when synthetic data proportions increased. For synthetic test data, results diverged. LDM_Conditional exhibited instability, with sharp drops in balanced accuracy and F1 scores across different training conditions. This suggests potential challenges in LDM-Conditional models when adapting to the nuances of synthetic data distribution. In contrast, DDPM-based models showed robust performance, maintaining high accuracy and F1 scores across mixed data training settings. These findings highlight DDPM’s efficacy in leveraging synthetic data for downstream tasks. The superior performance of DDPM_Conditional models compared to other configurations underscores the potential of conditional generative models in improving classification tasks using synthetic DTI data. One possible explanation is that the conditioning mechanism in DDPM_Conditional facilitates better alignment between the synthetic and real data distributions. This enhanced alignment likely aids the model in learning domain-invariant features crucial for sex classification. The poor performance of LDM_Conditional may be attributed to several factors, including suboptimal conditioning or artifacts in synthetic data generated by LDM. These results warrant further investigation into the representational fidelity of LDM-generated synthetic images, as discrepancies between synthetic and real distributions might lead to poor generalization.Interestingly, blending synthetic data with real data (e.g., 75% real + 25% synthetic) often yielded performance comparable to or exceeding 100% real data. This suggests that high-quality synthetic data can act as a data augmenter, introducing variability that prevents overfitting and enhances model robustness.

For the second downstream experiment, we evaluated the performance of CNN models for the same task of sex classification, but in the training dataset, real data is kept constant, and amount of synthetic data is increased in progressive runs by 25%. The balanced accuracy and F1 scores were assessed on both real and synthetic test datasets across four models: DDPM, DDPM_Conditional, LDM, and LDM_Conditional. In most cases, adding additional synthetic data while keeping the real data constant increases the balanced accuracies in the task. The DDPM reached accuracies of 0.98 when 50% additional Synthetic Data is added to the real training data in both tasks. This trend is echoed in F1 scores as well. Adding synthetic images from the LDM models also improved performance, but less in comparison to the DDPM models. The underperformance of LDM models reflects their limited ability to generalize between real and synthetic domains. This may stem from insufficient representation alignment or overfitting to domain-specific patterns. The experiments suggest that high-quality synthetic data can act as a data augmenter, increasing the accuracies of models.

## 5. CONCLUSION

In this paper, we tested the capability of various diffusion models to generate synthetic DTI-MD images. We evaluated strategies for training diffusion models effectively under the constraints of limited data, computational resources, and memory. Conditional latent diffusion models (LDM) and denoising diffusion probabilistic models (DDPM) were employed to generate synthetic 3D diffusion tensor imaging mean diffusivity (DTI-MD) MRI scans, with biological sex serving as a conditioning variable. Furthermore, we assessed the utility of these synthetic datasets in downstream applications, such as sex classification using convolutional neural networks, providing a practical benchmark for their relevance in data-driven neuroscience. Our findings underscore the potential of diffusion-based generative approaches to expand limited datasets and support various analytical pipelines. Future work could examine the added value of enhancing the diversity of synthetic data, optimizing model scalability for more extensive (or multi-site) datasets, and investigating additional downstream tasks, such as diagnostic classification of prognosis, to broaden their applications in biomedical research.

## Acknowledgements

This work was supported by NIH NIA grant U01 AG068057 (‘AI4AD’).

## Notes

### Competing Interest Statement

The authors have declared no competing interest.

### Summary of Updates

We have revised the following: 1) Updated upstream experiments with results for all models - DDPM, DDPM Conditional, LDM and LDM Conditional on 64x64x64 image size. 2) Updated with results from downstream sex classification tasks.

## References

[1] G. Kwon, et al. “Generation of 3D Brain MRI Using Auto-Encoding Generative Adversarial Networks,” Lect. Notes Comput. Sci. (including Subser. Lect. Notes Artif. Intell. Lect. Notes Bioinformatics), vol. 11766 LNCS, pp. 118–126, 2019.

[2] A.V. Dalca, et al. “Unsupervised learning of probabilistic diffeomorphic registration for images and surfaces.” Medical Image Analysis 57 (2019): 226–236.

[3] R. Rombach, et al., “High-Resolution Image Synthesis with Latent Diffusion Models,” in CVPR, 2022, pp. 10674–10685.

[4] J. Ho, et al., “Classifier-Free Diffusion Guidance.” NeurIPS 2021 Workshop on Deep Generative Models and Downstream Applications. 2021.

[5] W. H. L. Pinaya et al., “Brain Imaging Generation with Latent Diffusion Models,” in MICCAI workshop on Deep Generative Models (DGM4MICCAI), 2022, p. pp 117–126.

[6] A. Ijishakin, et al.”Interpretable Alzheimer’s Disease Classification Via a Contrastive Diffusion Autoencoder,” 2023, [Online]. Available: http://arxiv.org/abs/2306.03022.

[7] W. Peng et al., “Metadata-Conditioned Generative Models to Synthesize Anatomically-Plausible 3D Brain MRIs,” pp. 1–26, 2023, [Online]. Available: http://arxiv.org/abs/2310.04630.

[8] Z. Dorjsembe, et al. “Three-Dimensional Medical Image Synthesis with Denoising Diffusion Probabilistic Models,” in MIDL, 2022, pp. 2–4.

[9] W. Peng et al., (2023, October). ‘Generating realistic brain mris via a conditional diffusion probabilistic model.’ In International Conference on Medical Image Computing and Computer-Assisted Intervention (pp. 14–24). Cham: Springer Nature Switzerland.

[10] P. Sanchez, et al. “What is Healthy? Generative Counterfactual Diffusion for Lesion Localization,” Lect. Notes Comput. Sci. (including Subser. Lect. Notes Artif. Intell. Lect. Notes Bioinformatics), vol. 13609 LNCS, pp. 34–44, 2022.

[11] J. Wolleb, et al. “Diffusion Models for Medical Anomaly Detection,” Lect. Notes Comput. Sci. (including Subser. Lect. Notes Artif. Intell. Lect. Notes Bioinformatics), vol. 13438 LNCS, pp. 35–45, 2022.

[12] N.J. Dhinagar, et al. 2024. Counterfactual MRI Generation with Denoising Diffusion Models for Interpretable Alzheimer’s Disease Effect Detection. EMBC, 2024.

[13] Y. Li, et al. (2024). PASTA: Pathology-Aware MRI to PET Cross-Modal Translation with Diffusion Models. MICCAI. In International Conference on Medical Image Computing and Computer-Assisted Intervention (pp. 529–540). Cham: Springer Nature Switzerland.

[14] F. Deligianni, et al. (2016). NODDI and tensor-based microstructural indices as predictors of functional connectivity. PLoS One, 11(4), e0153404.

[15] E. Özarslan, et al. (2013). Mean apparent propagator (MAP) MRI: a novel diffusion imaging method for mapping tissue microstructure. NeuroImage, 78, 16–32.

[16] Y. Feng, et al. “Microstructural mapping of neural pathways in Alzheimer’s disease using macrostructure-informed normative tractometry.” Alzheimer’s Dement. 2024; 1–18.

[17] C. Owens-Walton, et al. (2024). ‘A worldwide study of white matter microstructural alterations in people living with Parkinson’s disease.’ npj Parkinson’s Disease, 10(1), 151.

[18] J. R. Taylor, et al. (2017). The Cambridge Centre for Ageing and Neuroscience (Cam-CAN) data repository: Structural and functional MRI, MEG, and cognitive data from a cross-sectional adult lifespan sample. NeuroImage, 144, 262–269.

[19] S. I. Thomopoulos, et al., “Diffusion MRI Metrics and their relation to Dementia Severity: Effect of Harmonization Approaches,” (2021) In 17th International Symposium on Medical Information Processing and Analysis (Vol. 12088, pp. 166–179). SPIE.

[20] T. Chattopadhyay, et al. (2023). Predicting dementia severity by merging anatomical and diffusion MRI with deep 3D convolutional neural networks. In the 18th International Symposium on Medical Information Processing and Analysis (SIPAIM; Vol. 12567, pp. 90–99). SPIE.

[21] A. Radford, et al.(2021, July). Learning transferable visual models from natural language supervision. In International conference on machine learning (pp. 8748–8763). PMLR.

[22] A. Gretton, et al., “A kernel two-sample test,” J. Mach. Learn. Res., vol. 13, pp. 723–773, 2012.

[23] D. Kingma, J. Ba (2015). “Adam: A method for Stochastic Optimization,” ICLR (2015).

[24] P. D. Tudosiu, et al. Realistic morphology-preserving generative modelling of the brain. Nat Mach Intell 6, 811–819 (2024).

[25] L. Puglisi, et al. (2024, October). Enhancing spatiotemporal disease progression models via latent diffusion and prior knowledge. In International Conference on Medical Image Computing and Computer-Assisted Intervention (pp. 173–183). Cham: Springer Nature Switzerland.

[26] B. van Breugel, et al. ‘Synthetic data in biomedicine via generative artificial intelligence.’ Nat Rev Bioeng 2, 991–1004 (2024).

